# A quality assurance framework in human intracranial electrophysiology

**DOI:** 10.64898/2026.08.06.743130

**Authors:** Noa Herz, Ruoyi Cao, Shiqi Qiu

## Abstract

Intracranial electroencephalography (iEEG) provides an unprecedented opportunity to directly record neural activity and causally perturb the human brain through electrical stimulation. Yet, the increasingly collaborative nature and complexity of modern iEEG studies pose substantial challenges for experimental control, data quality, and standardization. Unlike most experimental modalities, human iEEG data are acquired within dynamic clinical environments, where patient condition, recording quality, hardware configuration, and experimental protocols may vary across recording sessions and collaborating sites. The resulting heterogeneity creates opportunities for technical and procedural failures that often remain undetected until downstream analyses, when corrective action is no longer possible. Here, we present a framework for standardized session-level quality assurance in human iEEG research and provide an open-source implementation compatible with Brain Imaging Data Structure (BIDS)-organized datasets. The framework defines four complementary domains of quality assessment crucial for human iEEG studies: protocol fidelity, behavioral integrity, stimulation validation, and signal quality. These domains integrate electrophysiological recordings, behavioral event logs, and stimulation metadata to verify data completeness, confirm participant engagement, validate stimulation delivery, and identify potentially compromised recording channels. Automated quality metrics and standardized diagnostic visualizations are generated following each testing session, enabling rapid identification of technical and procedural failures while corrective action is still possible. By providing a standardized approach to session-level quality assurance, the framework improves data integrity, enhances reproducibility, facilitates analyst training, and supports harmonized data collection across laboratories and clinical sites.

## Introduction

Intracranial electroencephalography (iEEG) offers a rare opportunity to investigate complex cognitive processes in the human brain with a precision not achievable using noninvasive methods^1–3^. By combining millisecond-scale electrophysiological recordings with direct brain stimulation, iEEG has become a powerful platform for both mechanistic neuroscience and translational studies in psychiatric and neurological disease^4^. At the same time, iEEG studies pose unique technical, analytical, and logistical challenges. Unlike most noninvasive techniques, iEEG datasets are highly individualized, with the number, type, and placement of electrodes changing from patient to patient. In addition, data is often collected in complex hospital environments, where technical interruptions, clinical priorities, and hardware constraints can directly affect experimental execution. These conditions make iEEG studies vulnerable to consequential sources of error. As iEEG expands beyond its original clinical role to support increasingly collaborative and sophisticated neuroscience research, developing standardized quality-control tools will become crucial to ensuring transparent, reproducible, and harmonized data-acquisition practices.

Common challenges in iEEG research include malfunctioning or poorly referenced electrode contacts, line noise, incomplete or corrupted behavioral event logs, poor behavioral task performance, and stimulation artifacts that deviate from the intended temporal window or anatomical target. In multi-session studies, these challenges are often exacerbated, as errors can carry over across repeated data-collection time points. Despite the prevalence and potential impact of these sources of deviation, they are often identified only in later offline analyses, after the opportunity for remediation has passed, substantially reducing the scientific value of a rare and costly dataset. When issues are successfully recognized during data collection - primarily through visual inspection of ongoing iEEG recordings - their detection often depends on expert judgment that is time-consuming, subjective, and difficult to standardize across analysts, sessions, or research sites.

To address these problems, we present a quality control framework for iEEG research. While standardized quality assessment and preprocessing practices have been developed in the neuroimaging field (^5^ and ^6^), equivalent tools remain largely absent for human intracranial electrophysiology. Existing guidelines describe best practices for iEEG data acquisition, annotation, preprocessing, and sharing^7^, but little attention has been devoted to standardized quality-control procedures aimed at rapid detection of common experimental failures.

Here, we provide a principal method for rapid review of data collected in a single testing session, enabling the research team to identify common issues arising in human iEEG research. We provide a software implementation of this framework using an example dataset to demonstrate its application. By standardizing and automating these quality control procedures at the session level, this framework improves early error detection, reduces analyst burden, accelerates downstream analyses, and supports reproducible research practices across single and multi-site iEEG studies.

## Methods

### Framework Overview

The proposed quality assessment (QA) framework comprises four complementary domains that can be adapted to the requirements of different human intracranial electroencephalography (iEEG) experimental paradigms: **(1)** protocol fidelity, **(2)** behavioral integrity, **(3)** stimulation validation, and **(4)** signal quality (Figure 1). Together, these four domains provide a standardized approach for evaluating session-level data quality by integrating behavioral event logs, electrophysiological recordings, and stimulation metadata. Following testing of all QA domains, the framework automatically generates a standardized session-level report that includes quality metrics, detected deviations, and diagnostic visualizations, enabling rapid review after data acquisition. An example QA report and the source code for generating it are available in our lab’s public GitHub repository (https://github.com/HerzLab/QC-control-paper).

**Figure 1.**
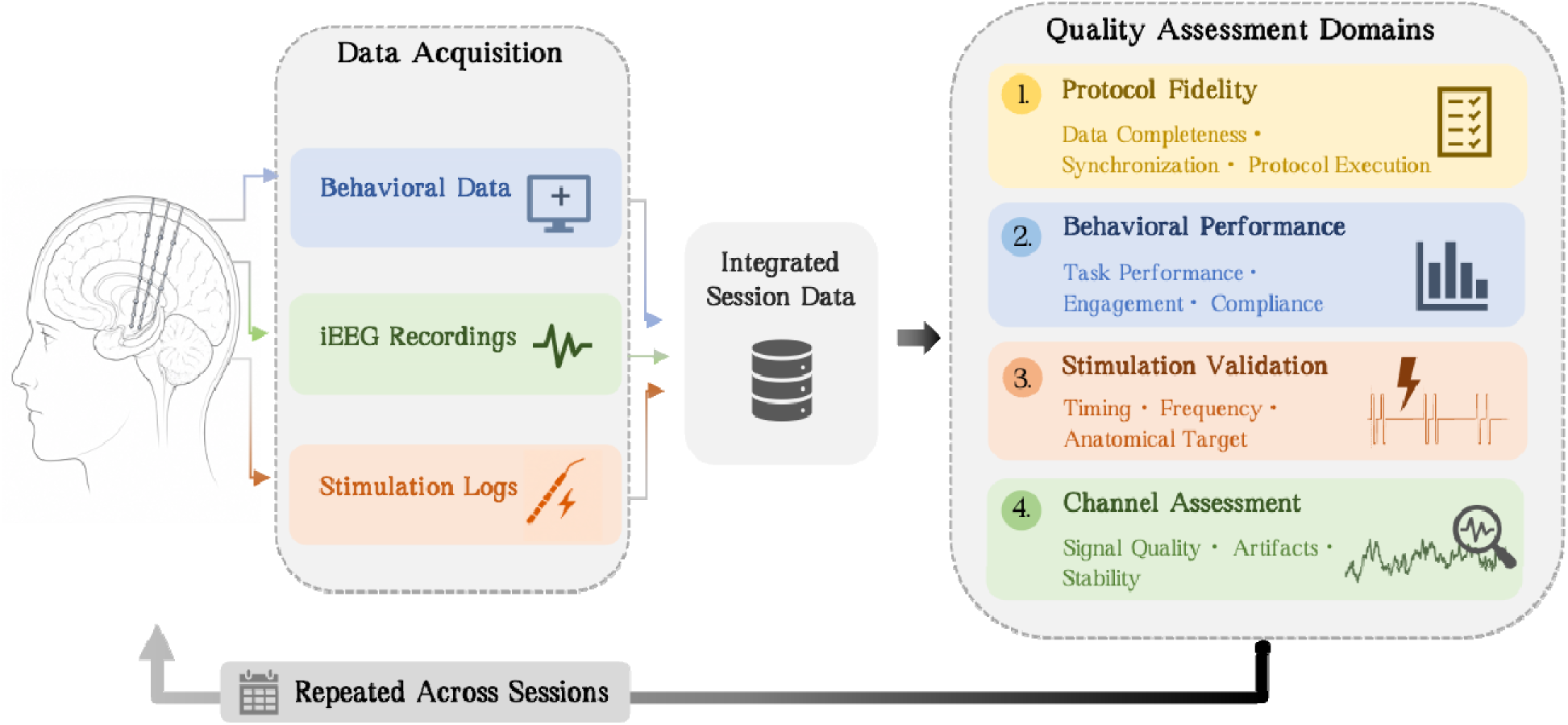
Schematics of the Quality Assurance Framework in Human Intracranial Electroencephalography.

### Example Application

To illustrate the framework, we apply it to an example iEEG data collected during a memory experiment involving direct electrical stimulation of the temporal lobe. This example dataset is available in BIDS format and can be downloaded from the Open Science Framework (https://osf.io/8yxde) to facilitate experimentation with and adoption of our report-generation framework.

The example dataset includes data from one epilepsy patient who performed a hybrid spatial-verbal memory task in which they acted as couriers navigating a virtual town. During the task’s encoding phase, the patient sequentially navigated to 12 stores. Upon arrival at each store, they were presented with the item delivered there (e.g., ‘Tomatoes’ at the Pizzeria). Upon the delivery of an item, a low-amplitude electrical stimulation (0.7 mA) was applied to predefined temporal lobe contacts for a 3-second duration, beginning at item presentation. Following completion of all 12 deliveries, the patient navigated to a final store before beginning a 90-second free-recall period, during which they verbally recalled as many delivered items as possible. Vocal responses were recorded and subsequently annotated offline. During retrieval, stimulation was delivered in alternating 3-second on/off trains using the same stimulation frequency assigned during encoding. Each session of this task included five trials, with stimulation frequency (3 or 8 Hz) counterbalanced across trials, permitting within-subject comparison of stimulation effects on memory encoding and retrieval.

Behavioral events were logged within the Unity task environment using millisecond-resolution timestamps. Stimulation commands, stimulation parameters, and delivery timestamps were recorded in parallel and synchronized with the electrophysiological recordings to produce a unified session-level dataset that served as input to the QA framework.

### Domain 1: Protocol Fidelity

The first quality assurance domain evaluates protocol fidelity by determining whether behavioral event logging and stimulation delivery faithfully reflect the intended experimental design. The primary objective of this domain is to identify missing data, synchronization failures, and procedural deviations before downstream behavioral or electrophysiological analyses are performed. Because many protocol errors invalidate an entire recording session regardless of signal quality, protocol fidelity serves as the first stage of session-level quality assurance.

To perform this assessment, behavioral event logs and electrophysiological/stimulation logs are extracted from their respective acquisition systems and merged into a unified, time-sorted data table using synchronized timestamps. Expected event types and event counts are defined *a priori* according to the experimental protocol. For each event type (e.g., *session_start* in Figure 1A), the observed number of events is compared with the expected count. In the example dataset presented here, each session is expected to contain five experimental trials (Figure 1A). Events are classified as *PASS* when observed counts match protocol-defined expectations and *FAIL* when discrepancies are detected. Event-count discrepancies typically reflect incomplete data acquisition, incorrect task implementation, failures in event transmission between software components, or synchronization problems between acquisition systems. Sessions failing critical protocol checks are flagged for manual review before subsequent quality assurance procedures or scientific analyses are performed.

In studies involving direct electrical brain stimulation, verification of stimulation delivery represents an additional component of protocol fidelity. Deviations from the intended stimulation parameters, incorrect electrode targets, or stimulation delivered outside the assigned task phase may substantially alter both neural responses and behavioral outcomes. Accordingly, stimulation events are grouped according to their commanded parameters and experimental phase (Figure 1B). For each trial, the framework verifies the expected number of stimulation pulses and summarizes stimulation delivery according to task phase, stimulation frequency, and stimulation target. This structured summary enables rapid detection of missing stimulation trains, incorrect task-phase assignments, unexpected stimulation frequencies, or stimulation delivered to unintended electrode contacts. Such deviations most commonly arise from errors in stimulation program configuration, event-triggering logic, or communication between the behavioral software and the stimulation control hardware during data acquisition.

### Domain 2: Behavioral Integrity

The second quality assurance domain evaluates behavioral integrity by determining whether participant performance is consistent with successful task completion and meaningful interpretation of the recorded neural data. Unlike formal behavioral analyses, which are designed to test scientific hypotheses, this module aims to identify deviations that may indicate incomplete data acquisition, poor task comprehension, disrupted engagement, or other session-level issues that compromise data quality. Behavioral quality assurance, therefore, focuses on verifying that the experimental conditions required for a valid interpretation of the electrophysiological recordings were met. Behavioral analyses that do not directly inform these quality-control goals are not only redundant at this stage, but may also introduce bias by encouraging premature interpretation of outcomes or implicitly influencing data exclusion decisions.

To support this assessment, the pipeline generates standardized summaries of task performance tailored to the experimental paradigm. In the example dataset presented here, recall performance serves as a principal indicator of session integrity because successful recall depends on multiple components of the recording workflow, including proper task execution, successful synchronization of behavioral events, accurate recording of verbal responses, and sustained participant engagement (Figure 2). Session-level summaries include recall accuracy in each trial, overall recall performance, and intrusion (false recall) rates. Abrupt declines in recall performance, unusually high intrusion rates, or atypical response distributions may indicate interruptions during testing, microphone failures, misunderstanding of task instructions, or reduced participant engagement.

**Figure 2.**
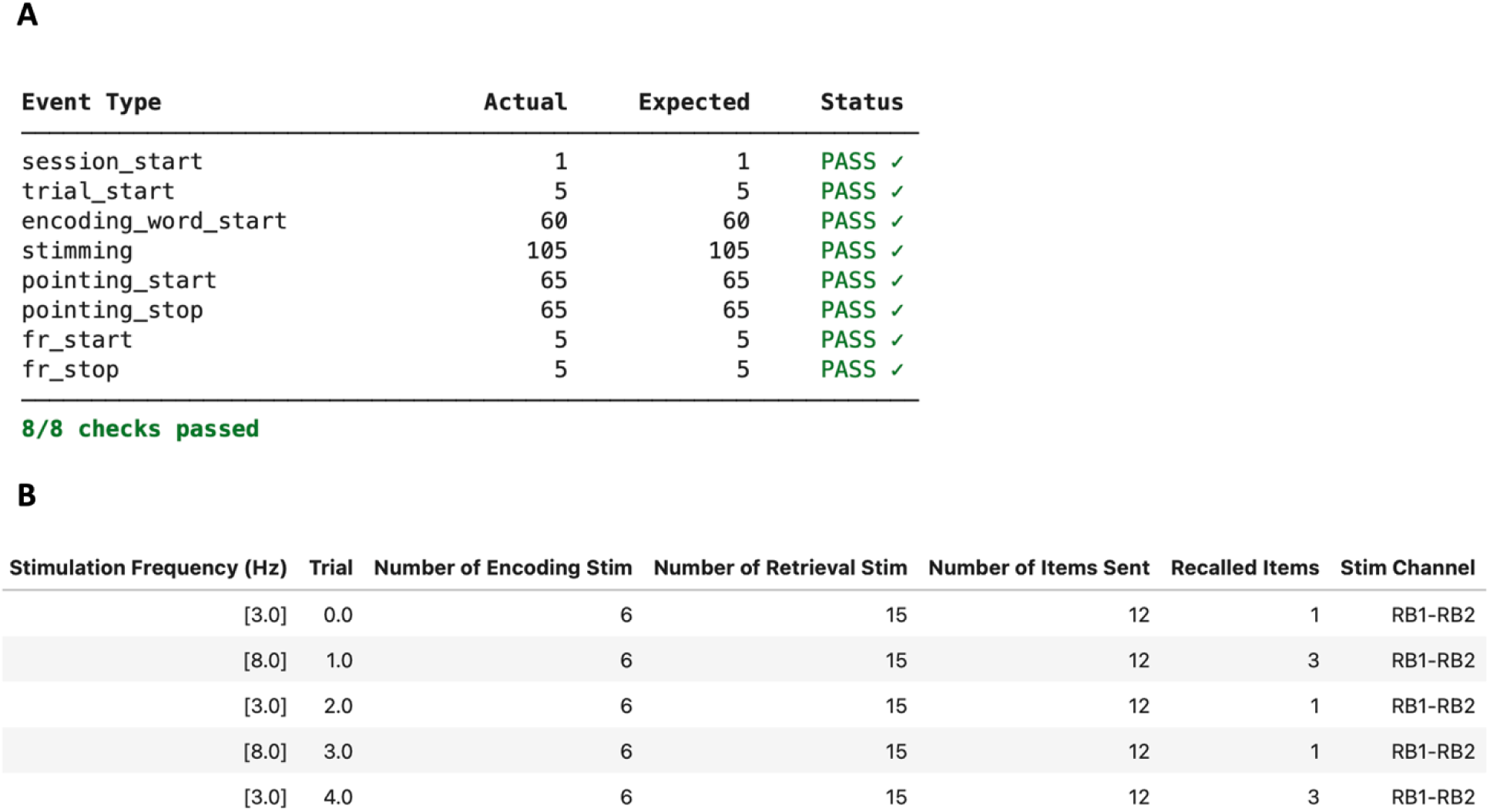
Example protocol adherence summary for a representative iEEG session. **A. Event-count verification table.** “Event Type” describes all conditions included in the cognitive task, as defined by the experimental program. In this example dataset, each session begins with a “session_start” notice, and includes five trials (marked here by “trial_start”). All events during the testing session should appear in the table above to ensure proper task administration. **B**. Example stimulation categorization report for a single session comprising five trials. Session-level summaries of stimulation events are stratified by task phase (here, encoding vs. retrieval) and include stimulation frequency (alternating between 3 and 8 Hz) and stimulation target (here, a single bipolar electrode pair labeled RB1–RB2). This structured summary enables rapid verification of protocol adherence and facilitates identification of deviations from intended stimulation parameters.

**Figure 2.**
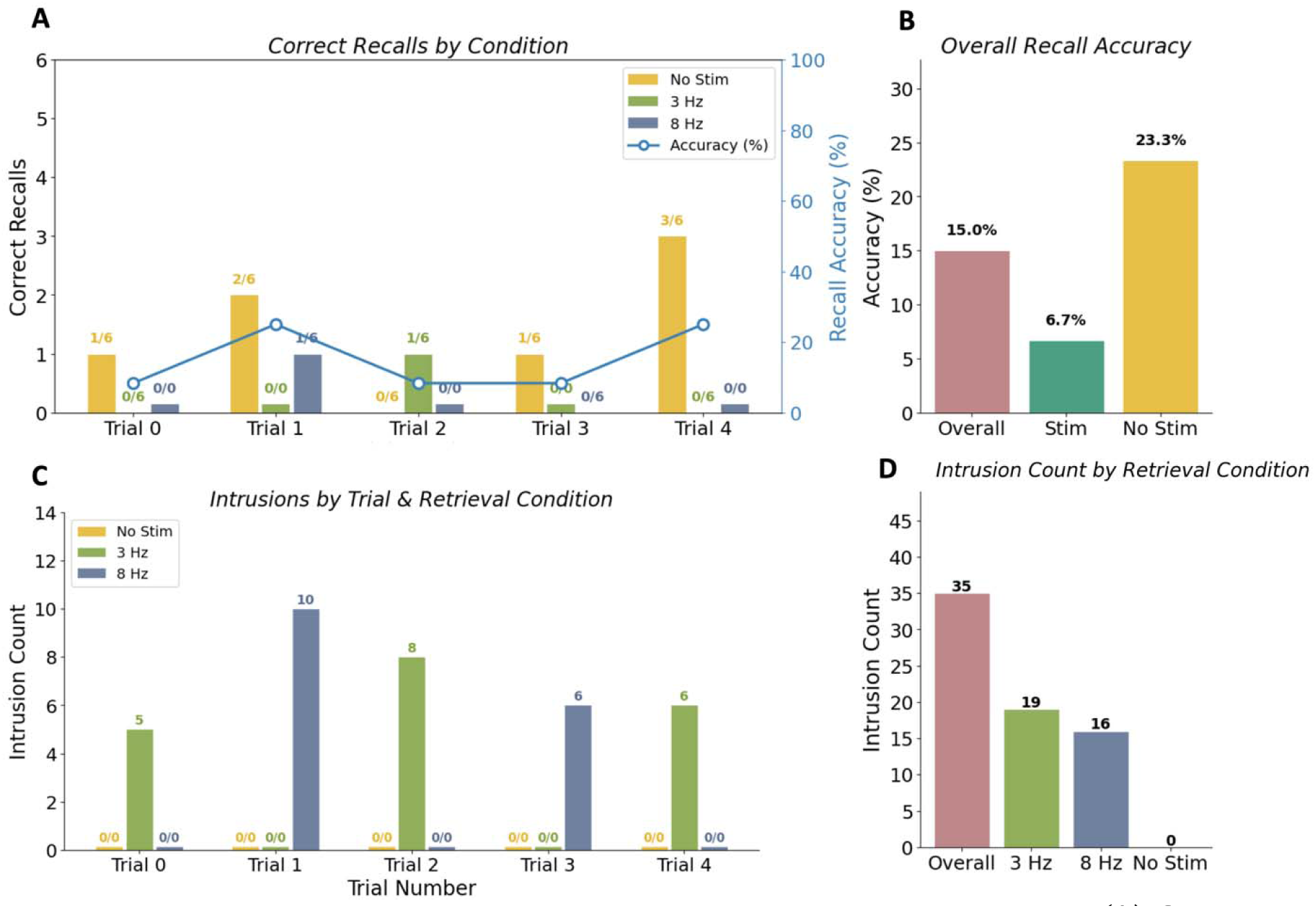
Recall performance summaries for session-level quality assessment. **(A)** Correct recalls per trial (left y-axis, bars) and overall recall accuracy (right y-axis, blue line) across the five experimental trials (Trials 0–4). Abrupt memory declines may reflect session interruptions or inconsistent logging, whereas gradual performance drift may indicate fatigue or reduced engagement. **(B)** Overall recall accuracy collapsed across trials. **(C, D)** Intrusion counts by retrieval condition (No Stim - yellow; 3 Hz - green; 8 Hz - blue), shown per trial **(C)** and collapsed across trials **(D).** Unusually low accuracy or elevated error rates may warrant further inspection for potential misunderstanding of task instructions or failure to record participant input.

Additional behavioral measures, including navigation times and spatial trajectories (Figure S1), provide complementary indicators of task performance. Prolonged navigation times, restricted movement patterns, or incomplete exploration of the virtual environment may suggest joystick malfunction, fatigue, or difficulty interacting with the task.

Although the present example focuses on a spatial-verbal memory task, quality assurance metrics may instead include reaction times, recognition accuracy, confidence ratings, motor responses, language production, eye-tracking measures, or other task-specific behavioral variables. The selected metrics should be chosen according to the scientific paradigm and the anticipated sources of session failure.

### Domain 3: Stimulation Validation

The third quality-assurance domain evaluates whether electrical stimulation was delivered in accordance with the intended experimental protocol. Since stimulation serves as the experimental manipulation in many iEEG studies, deviations in stimulation timing, temporal structure, or anatomical targeting can fundamentally alter both neural and behavioral outcomes. Verification of stimulation delivery is therefore essential for interpreting the experiment’s results and for ensuring adherence to stimulation safety protocols.

Stimulation validation comprises three complementary components: Stimulation timing, parameters, and anatomical specificity. Together, these assessments determine whether stimulation occurred at the intended time point relative to the cognitive task, with the programmed stimulation parameters, and at the intended anatomical location. Rather than relying solely on stimulation-command logs, the framework performs these assessments directly from the recorded electrophysiological signal, providing independent confirmation of actual stimulation delivery.

For these purposes, the electrophysiological signal during stimulation is evaluated against the intended parameters of the stimulation protocol. This alignment enables detection of deviations such as incorrect stimulation frequency, timing offsets relative to behavioral events, incomplete or truncated stimulation trains, and unintended propagation of stimulation artifacts to neighboring electrodes. The module implements signal-level analyses that include (i) alignment of raw voltage traces to declared stimulation onsets, (ii) generation of stimulation-locked averages to assess temporal precision and frequency fidelity, and (iii) characterization of stimulation artifacts across electrode contacts to assess spatial specificity. Together, these analyses enable rapid identification of timing offsets, incorrect stimulation parameters, incomplete or truncated stimulation trains, and unintended spread of stimulation artifacts, providing a comprehensive assessment of stimulation delivery before downstream analyses.

#### Stimulation Timing

Precise temporal alignment between electrical stimulation and behavioral task events is essential for studies investigating the causal effects of stimulation on cognition. Protocols that deliver stimulation contingent on task or behavioral events require accurate synchronization between stimulation onset and predefined task conditions. While the specific synchronization mechanisms depend on the acquisition hardware and software, verification of stimulation timing is a necessary step to ensure experimental validity.

To assess timing accuracy, raw voltage traces are aligned with the declared stimulation onset times obtained from the experimental metadata. Single-trial overlays are generated, with stimulation epochs explicitly marked (Figure 3, top). This approach enables detection of systematic timing offsets, trial-to-trial variability in onset latency, or complete failures of stimulation delivery. To further evaluate temporal precision, a zoomed-in view of the onset period is examined (here, −20 to 50 ms relative to declared stimulation onset; Figure 3, bottom). This fine temporal window allows detection of subtle misalignments between the logged stimulation time and the actual onset of stimulation artifacts in the electrophysiological signal.

**Figure 3.**
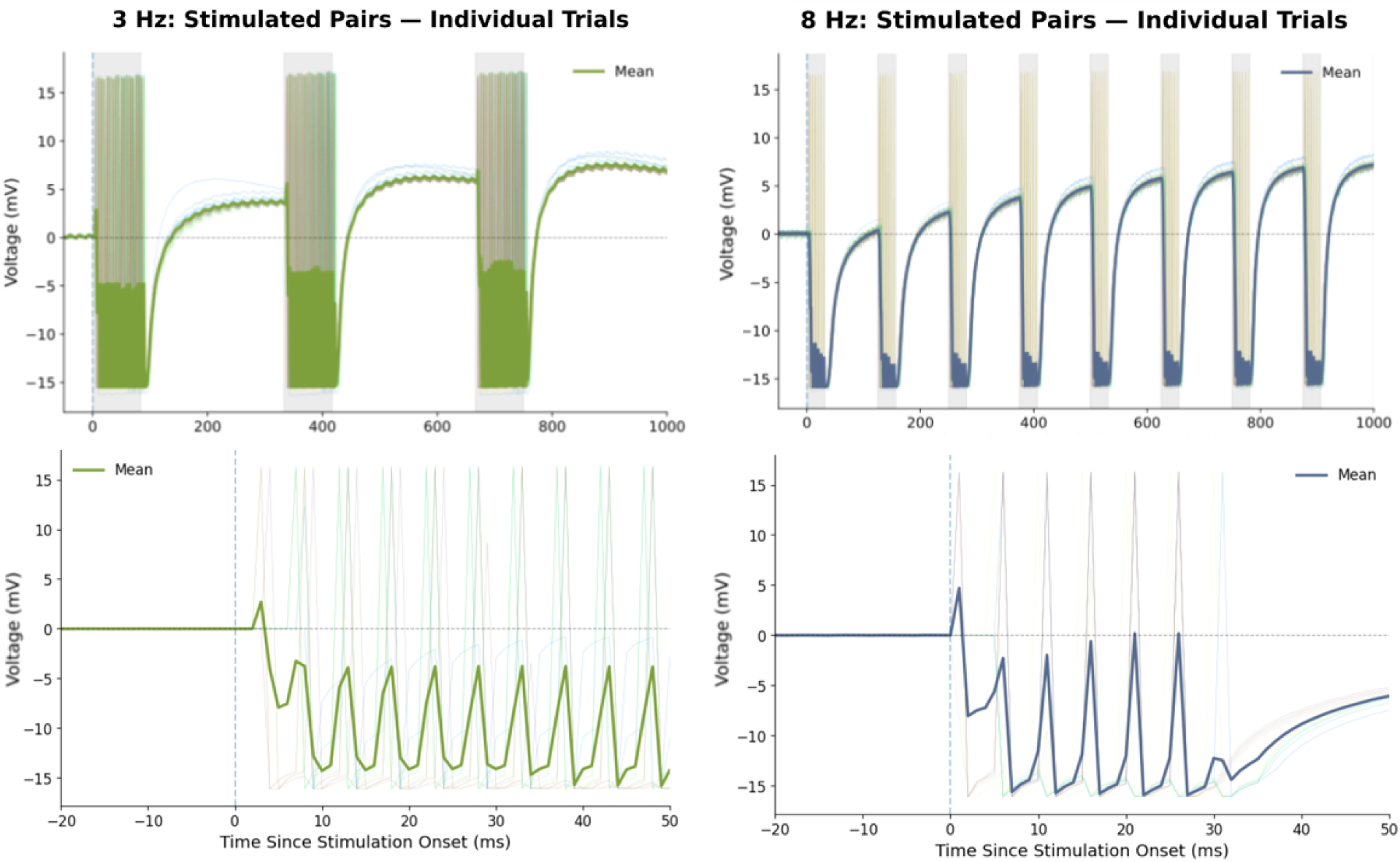
Verification of stimulation timing in iEEG recordings. Individual iEEG traces from stimulated contact pairs are aligned to the stimulation-onset timestamps from the task event logs. Trial-averaged responses are overlaid in green (3 Hz) and blue (8 Hz). **Top:** 1000-ms epochs for 3 Hz (left) and 8 Hz (right) stimulation delivery. Stimulation artifacts recur at the expected inter-burst intervals (∼333 ms and ∼125 ms, respectively), with burst durations of ∼83 ms and ∼31 ms (25% duty cycle), confirming delivery at the intended stimulation frequencies and duty cycle across trials. **Bottom:** Expanded stimulation-onset view (−20 to +50 ms) for 3 Hz (left) and 8 Hz (right) stimulation delivery. The dashed vertical line at 0 denotes the stimulation onset timestamp derived from the event logs. The first stimulation artifact is consistently aligned to the onset marker across trials, with artifact onset occurring within ∼5 ms of the logged timestamp, indicating accurate synchronization between the stimulation device and the behavioral task recording system.

At the single-trial level, consistent alignment of stimulation artifacts across trials provides evidence of reliable timing, whereas variability in onset latency suggests instability in stimulation delivery or timing jitter in the synchronization pipeline. Trials in which no stimulation artifact is observed despite a logged event are indicative of stimulation delivery failure. Deviations beyond the expected range of temporal variability may indicate synchronization errors between acquisition systems, delays in trigger transmission, or inaccuracies in event logging.

#### Stimulation Frequency and Spatial Specificity

In addition to timing accuracy, it is essential to confirm that stimulation is delivered with the intended temporal structure (frequency) and spatial confinement. Early detection of protocol deviations is critical for both experimental validity and patient safety. Deviations such as prolonged trains, elevated amplitudes or frequencies, or stimulation reaching unintended contacts near suspected seizure-onset zones may exceed safety limits and increase the risk of after-discharges or seizure provocation.

Spatial specificity is assessed by characterizing the distribution of stimulation artifacts across contacts. Electrical stimulation should produce the largest artifacts at the stimulated contacts, with attenuated effects on adjacent contacts along the same electrode shank and minimal or absent artifacts on more distant electrodes. Deviations from this pattern may indicate an incorrect stimulation configuration, referencing issues, or hardware malfunction. To systematically evaluate both spatial specificity and stimulation frequency, electrodes are grouped into three configurations (Figure 4): (A) Stimulated contacts; used to verify stimulation timing and confirm the expected pulse frequency; (B) Non-stimulated contacts on the stimulated shank; used to assess the magnitude of local propagation of stimulation artifacts; and (C) Contacts on other, non-stimulated, shanks; used to verify minimal distal contamination. For each configuration, stimulation-locked responses are computed and visualized. In the stimulated contact pair (Figure 4A), regular, periodic deflections corresponding to the programmed stimulation frequency (Here, 3 Hz or 8 Hz) confirm the correct temporal structure of stimulation delivery. Attenuation of these deflections in same-shank non-stimulated contacts (Figure 4B), and their near absence in distant contacts (Figure 4C), supports appropriate spatial confinement.

**Figure 4.**
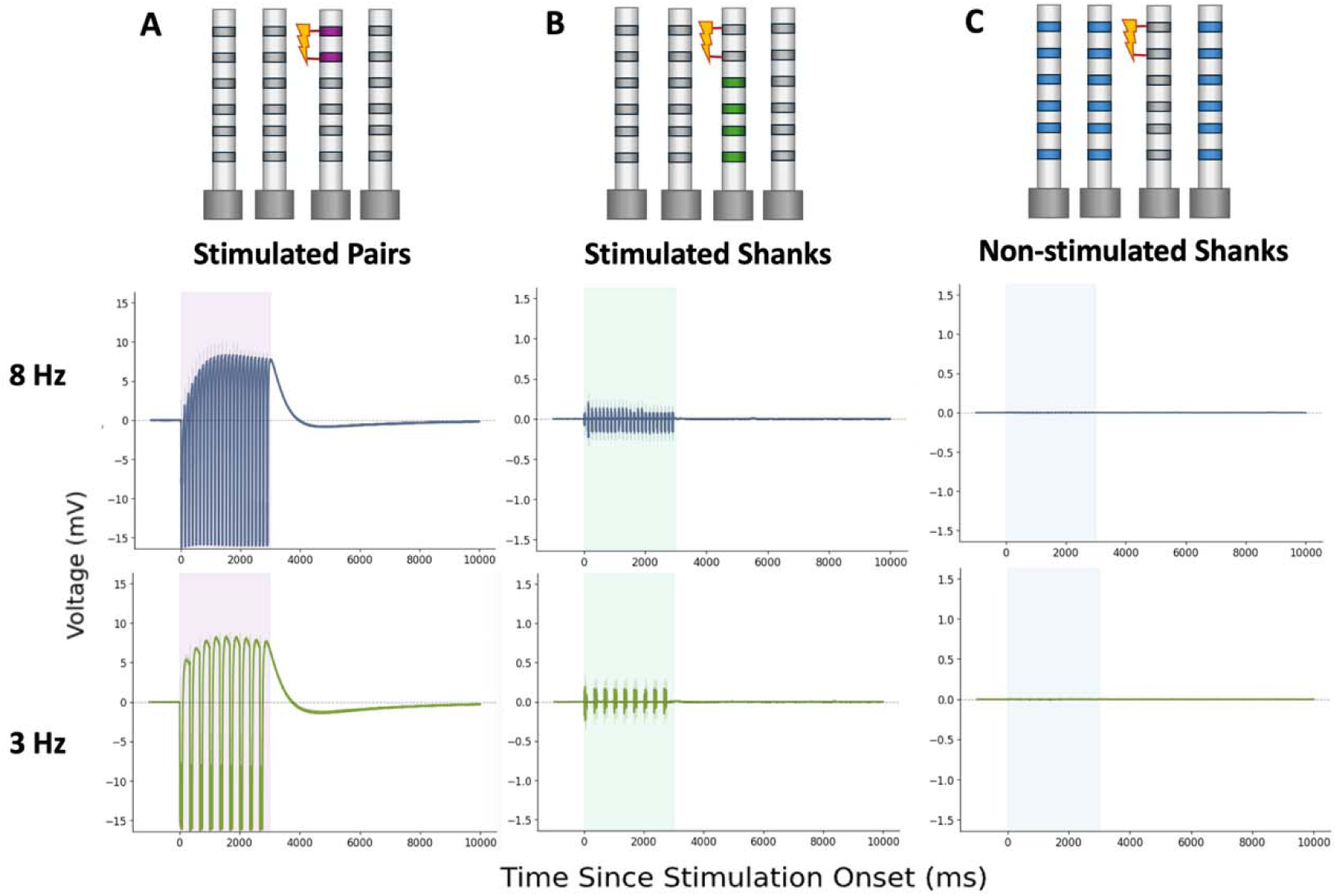
Electrode-contact grouping to assess stimulation spatial specificity. Electrode contacts are partitioned into three configurations, illustrated schematically at the top of each column. Each configuration shows the averaged signal in: **(A)** stimulated contacts; **(B)** non-stimulated contacts on the stimulated shank; and **(C**) contacts on non-stimulated shanks. Stimulus-locked averages are shown here for 8 Hz (blue) and 3 Hz (green) stimulation. The shaded region denotes the 3-second stimulation interval. Artifact amplitude is maximal at the stimulated pair, attenuated on same-shank non-stimulated contacts, and minimal on distant shanks, demonstrating the expected spatial decay of stimulation-related artifacts (note that the y-axis range in panels B and C is scaled to one-tenth of panel A).

This structured comparison enables rapid detection of stimulation errors, prompting the research team to inspect the affected session, verify hardware and referencing configurations, and, when needed, adjust stimulation parameters or the experimental setup in subsequent recording sessions.

### Domain 4: Signal Quality

The fourth quality-assurance domain evaluates whether recorded electrophysiological signals are of sufficient quality for reliable scientific interpretation. Although intracranial EEG is substantially less susceptible than scalp EEG to physiological artifacts such as eye movements, muscle activity, and cardiac activity, iEEG recordings remain vulnerable to electrical contamination, unstable electrode contacts, hardware malfunction, inappropriate referencing, stimulation-related artifacts, and other sources of signal degradation that can compromise downstream analyses.

During data collection, trained experimenters routinely inspect raw signals to identify recording problems. However, manual inspection becomes increasingly difficult as electrode coverage expands to hundreds of implanted contacts and recordings accumulate across multiple sessions. Moreover, the dynamic hospital environment introduces time-varying noise sources arising from changes in patient position, electrode impedance, referencing configuration, nearby electronic equipment, and clinical activity. These sources of signal degradation may affect only subsets of channels or emerge intermittently across recording sessions, making systematic assessment of signal quality both challenging and essential. Automated post-session quality assurance therefore provides a complementary layer of validation beyond online signal inspection by systematically identifying channels requiring expert review.

The present framework evaluates signal quality using complementary spectral and statistical measures that capture distinct forms of signal degradation. Spectral analyses identify frequency-specific abnormalities indicative of electrical contamination or recording-system malfunctions, whereas statistical measures quantify signal instability and other deviations from the expected characteristics of neural recordings. Rather than automatically excluding channels, these measures identify recordings that warrant further inspection and provide standardized visualizations to support transparent and reproducible quality-assurance decisions.

#### Spectral Assessment

Transforming electrophysiological signals from the time domain into the frequency domain provides a powerful means of characterizing recording quality. Whereas time-domain signals describe voltage fluctuations over time, spectral decomposition quantifies how signal power is distributed across frequencies, enabling frequency-specific sources of contamination to be readily identified. Spectral analyses therefore complement visual inspection of raw traces by revealing abnormalities that may be difficult to detect by eye through looking at the time domain.

Physiological neural recordings typically exhibit a 1/f spectral profile (pink noise), in which signal power decreases gradually with increasing frequency (Figure 5). Marked deviations from this expected spectral shape, including unusually flat spectra or unexpected narrow-band peaks, may indicate recording artifacts, amplifier saturation, poor electrode contact, or other technical abnormalities.

**Figure 5.**
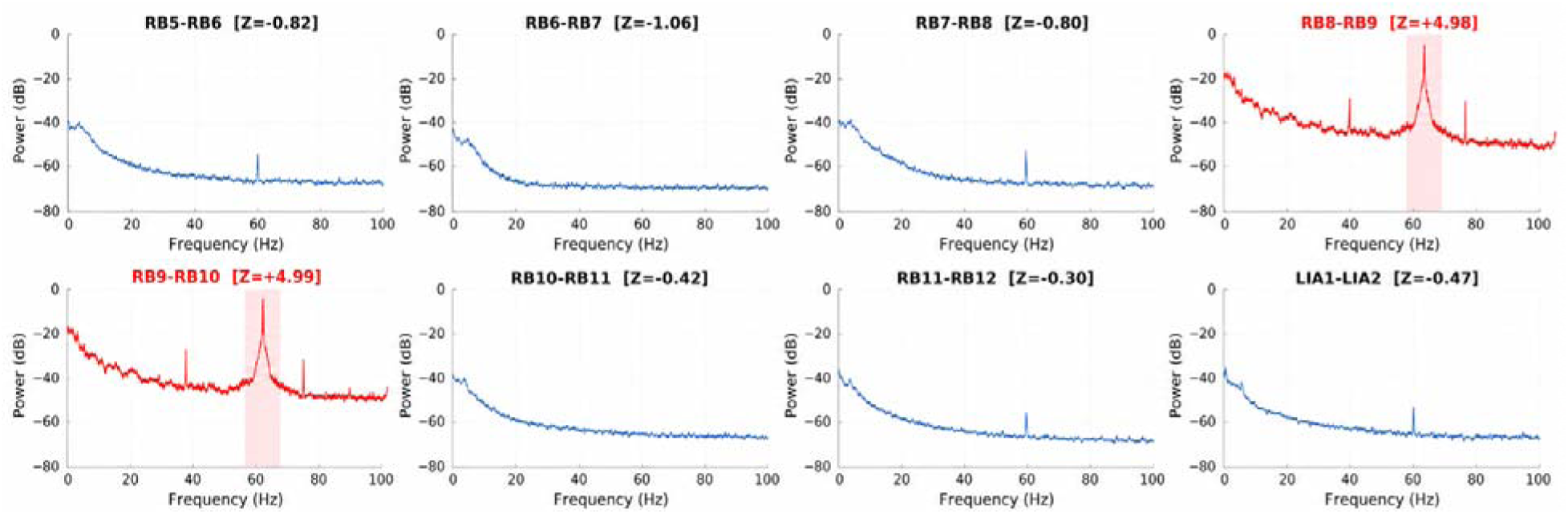
Spectral assessment of line-noise contamination. Log-transformed power spectra (dB) are shown for 8 example bipolar channel pairs across the 0–100 Hz frequency range. Spectra were computed by applying a fast Fourier transform (FFT) to single-trial iEEG signals for each contact, followed by averaging across trials. Channels whose mean power within the 55–65 Hz frequency band exceeded 2 standard deviations above the session mean are automatically highlighted in red, with the 55–65 Hz frequency band shaded; all remaining channels are shown in blue. In this example, contacts RB8–RB9 and RB9-RB10 exhibited elevated 60 Hz spectral peaks and were therefore automatically flagged for review.

Line noise serves as a particularly informative example of spectral contamination. Line noise arises from the mains electricity supply and appears as a narrow-band oscillation at 60 Hz in the United States (50 Hz in many other countries). Although line noise can often be attenuated during preprocessing using notch filtering or spectral regression, excessive line-noise power frequently reflects broader problems with signal acquisition, including poor grounding or referencing, elevated electrode impedance, hardware malfunction, or environmental electromagnetic interference. Consequently, line noise functions not only as a source of signal contamination but also as a useful diagnostic indicator of recording-system integrity. These underlying issues may degrade signal quality well beyond the narrow frequency band of the electrical mains and can substantially affect downstream analyses, particularly those examining high-frequency neural activity.

### Line Noise Detection

Because line noise occupies a well-defined frequency band, it can be readily identified after transforming the signal to the frequency domain via spectral decomposition (implemented here using the fast Fourier transform, FFT). In the present implementation, mean power within the 55–65 Hz band is computed for each channel and standardized across channels within a recording session using z-scores. Power spectra for all channels are displayed, with channels exceeding a predefined threshold (z > 2) automatically highlighted for visual inspection (Figure 5 and Figure S2).

Because line-noise power is standardized within each recording session, the resulting z-scores characterize the relative distribution of contamination across channels rather than its absolute magnitude. This relative representation facilitates identification of spatial patterns that often provide greater diagnostic value than the absolute level of line-noise power. To visualize these patterns, the framework generates a scatter plot showing each channel’s mean mains-frequency power while preserving the spatial organization of the implanted electrodes (Figure 6). Channels are arranged according to standard clinical electrode nomenclature, enabling rapid identification of electrode clusters. Elevated line-noise power confined to a single contact may indicate localized contact problems, elevated impedance, or contact-specific contamination. In contrast, diffuse elevations spanning multiple electrodes are more suggestive of referencing errors, grounding problems, or environmental electrical interference.

**Figure 6.**
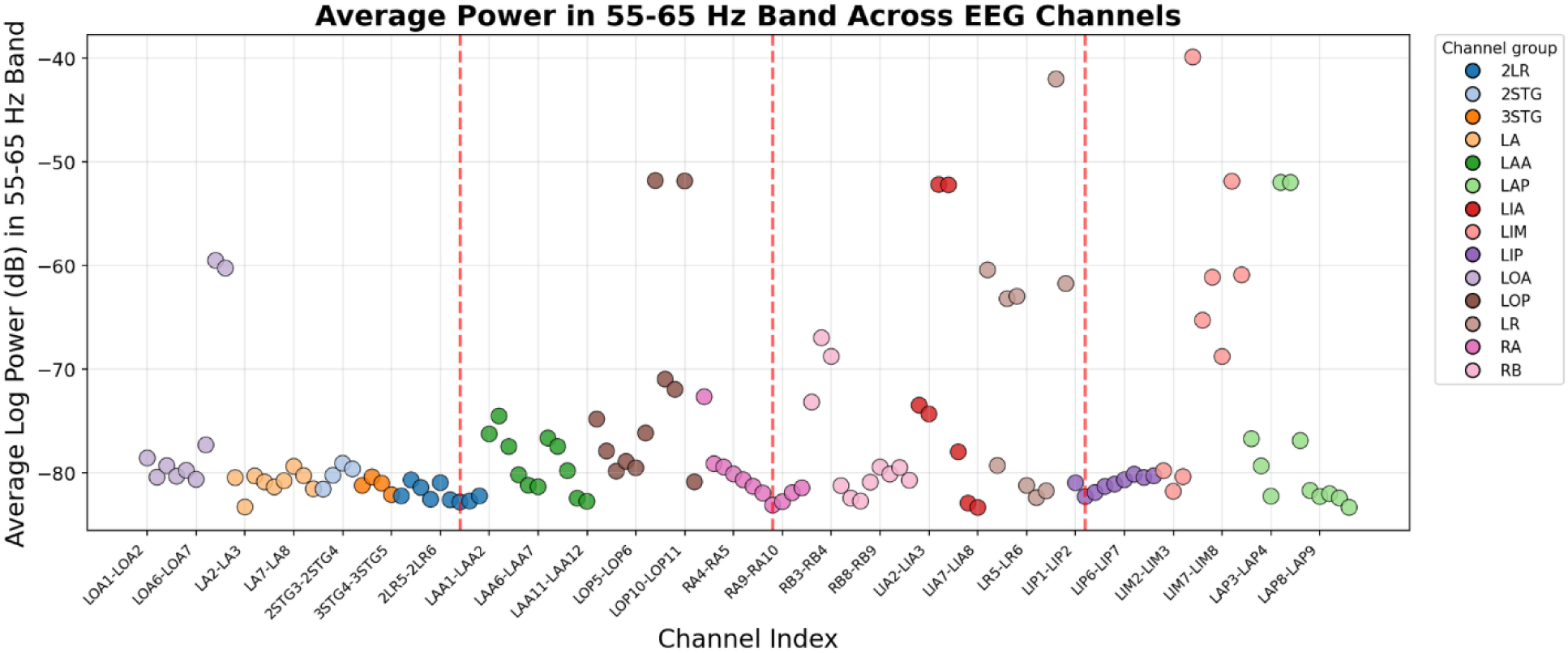
Spatial assessment of line-noise contamination. Each point represents a single bipolar channel pair and shows the mean log-transformed power (dB) in the 55–65 Hz band. Channels are grouped along the x-axis by clinical electrode lead names (e.g., LOA, LA) to facilitate visualization of spatial clustering among implanted electrodes. Vertical red dashed lines demarcate the boundaries between recording banks, which, in the present recording system, include 32 contacts. Spatial clustering of elevated line-noise power confined to a single contact may indicate localized contact problems, elevated impedance, or electrode-specific contamination. In contrast, clustering across channels sharing the same acquisition bank may suggest connector contamination, compromised shielding, or other hardware-specific noise sources affecting multiple contacts simultaneously.

Interpretation of these spatial patterns can be further refined by considering the recording-system architecture. Channels sharing amplifier ports, headstages, or connector banks may exhibit common noise profiles if contamination arises from hardware-specific failures. Consequently, integrating the spatial distribution of line noise with knowledge of the acquisition system provides valuable insight into the underlying source of recording artifacts and can guide efficient troubleshooting during ongoing data collection.

#### Channel Quality Assessment Using Statistical Variability

Unlike line noise, which appears as a relatively stable oscillatory signal at a fixed frequency, many recording artifacts are transient, irregular, and difficult to identify using spectral analyses alone. Instead of producing contamination within a narrow frequency band, these artifacts often alter the statistical properties of neural recordings, resulting in abnormal signal variability across channels, trials, or time. Statistical characterization of signal variability therefore provides a complementary approach for identifying recording abnormalities that may not be observable in the frequency domain.

Reliable neural recordings are expected to exhibit relatively stable statistical properties both across repeated trials and within individual trials over time. Excessive variability across trials may reflect intermittent noise sources, unstable electrode contacts, inconsistent referencing, transient movement-related disturbances, or trial-specific recording artifacts. Conversely, excessive variability within individual trials may indicate persistent broadband noise, slow voltage drift, amplifier saturation, or residual stimulation artifacts.

To capture these complementary forms of instability, the framework quantifies trial-to-trial and temporal variability. Trial-to-trial variability measures the consistency of neural responses across repeated observations, identifying channels whose responses vary abnormally between trials despite exhibiting stable temporal dynamics within each trial. In contrast, temporal variability quantifies signal fluctuations over time within individual trials, identifying channels with excessive temporal instability even when these fluctuations are highly reproducible across trials.

Importantly, these metrics characterize distinct aspects of signal quality. For example, channels exhibiting robust event-related potentials (ERPs) may show substantial temporal variability due to large physiological deflections, yet demonstrate low trial-to-trial variability because the response is reproducible across trials. In such cases, elevated temporal variance reflects genuine neural activity rather than recording instability. Conversely, artifacts arising from unstable electrodes or intermittent recording failures frequently increase both trial-to-trial variability and temporal variance. Considering these measures jointly provides complementary perspectives on channel integrity and facilitates interpretation of the underlying sources of signal abnormality.

For both the trial-to-trial and temporal metrics, variability values are standardized across channels using z-scores, and channels exceeding predefined thresholds are flagged for review. The framework generates standardized visualizations - including variability heatmaps, channel-level summary statistics, and representative waveforms from flagged channels (Figures 7 and 8) - to facilitate rapid identification of atypical recording behavior and promote reproducible quality assessment across sessions.

**Figure 7.**
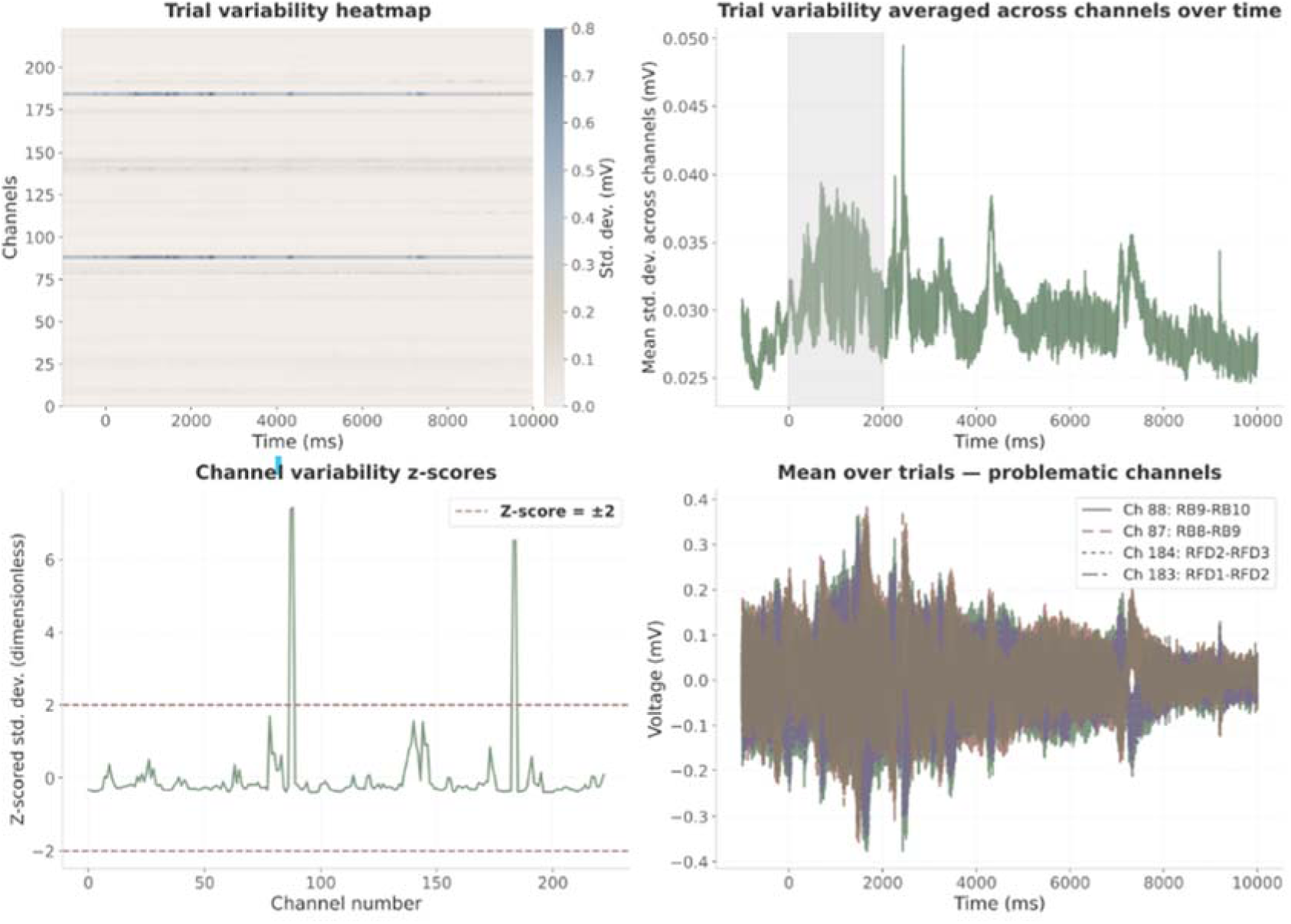
Signal Quality Assessment via Trial-to-Trial Variability. **Upper left:** Heatmap showing standard deviation across trials for each channel (y-axis) over an example 10-second recording epoch (x-axis). This visualization enables rapid identification of channels with persistently elevated variability and allows assessment of whether instability is sustained throughout the epoch or confined to specific temporal windows. **Upper right:** Mean trial-to-trial variability across all channels as a function of time. The item-presentation period is highlighted in gray to distinguish task-related signal fluctuations from background noise occurring outside the primary period of interest. Peaks restricted to task periods may reflect physiological event-related responses, whereas elevated variability throughout the epoch is more suggestive of nonphysiological artifact. **Bottom left:** Z-scored channel variability metrics for all channels. Horizontal dashed red lines indicate the predefined ±2 z-score threshold used to flag potentially problematic channels. Channels exceeding this threshold exhibit abnormally high or low trial-to-trial consistency relative to the session-wide distribution. Anatomical grouping of flagged channels also facilitates identification of spatial clustering, such as multiple noisy contacts located on the same electrode shank, which may suggest shared reference contamination, poor electrode contact, or localized hardware instability rather than an isolated single-channel artifact. **Bottom right:** Trial-averaged waveforms from channels with the highest absolute variability z-scores. These channels exhibit the greatest inconsistency across repeated trials and therefore warrant targeted visual inspection.

**Figure 8.**
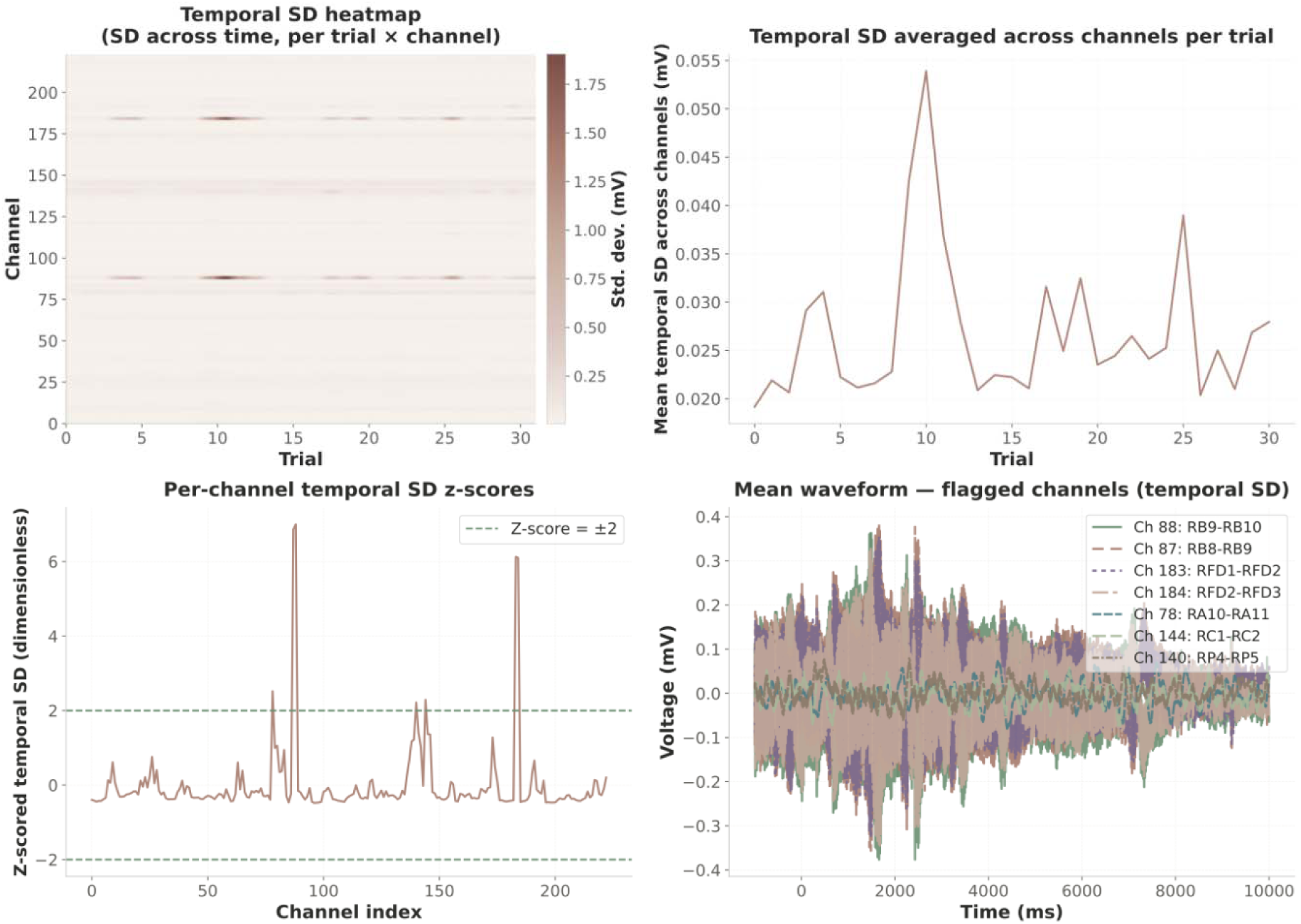
Signal Quality Assessment via Temporal Variability. **Upper left:** Heatmap showing the standard deviation for each channel (y-axis) and trial (x-axis). Darker colors indicate greater within-trial temporal variability. Persistently dark rows identify channels with consistently unstable signals, whereas dark columns may indicate trials contaminated by broadband noise, movement-related artifact, or transient recording instability. **Upper right:** Mean temporal variance averaged across channels as a function of trial. Gradual monotonic changes across trials may indicate slow signal drift over the course of the experiment, whereas exceptionally high variance confined to a small number of trials may suggest transient global contamination affecting those trials. **Bottom left:** Z-scored temporal variability metric for each channel. Horizontal dashed lines indicate the predefined ±2 z-score thresholds used to flag potentially problematic channels. Channels exceeding these bounds exhibit abnormally high or low temporal variability relative to the session-wide distribution. Clustering of flagged channels on the same electrode shank may indicate a shared reference problem, poor electrode contact, or hardware instability rather than isolated single-channel artifact. **Bottom right:** Trial-averaged waveforms for the channels with the highest absolute temporal variability z-scores. These channels exhibit excessive within-trial instability and warrant further review.

Because statistical metrics alone cannot fully distinguish physiological variability from recording artifacts, automated assessments are complemented by direct visual inspection of the underlying electrophysiological signals. To support this final review step, the framework generates overview plots displaying raw waveforms from all recorded channels simultaneously (Figure 9), with channels flagged by either variability metric highlighted for immediate inspection. The proposed metrics are not intended as automated channel-rejection criteria. Decisions regarding channel exclusion depend on multiple factors, including the experimental paradigm, analysis objectives, anatomical location, and the nature of the observed abnormality. Instead, these statistical measures serve as practical screening tools that prioritize channels for expert review, enabling more efficient and standardized quality assurance during data collection and preprocessing.

**Figure 9.**
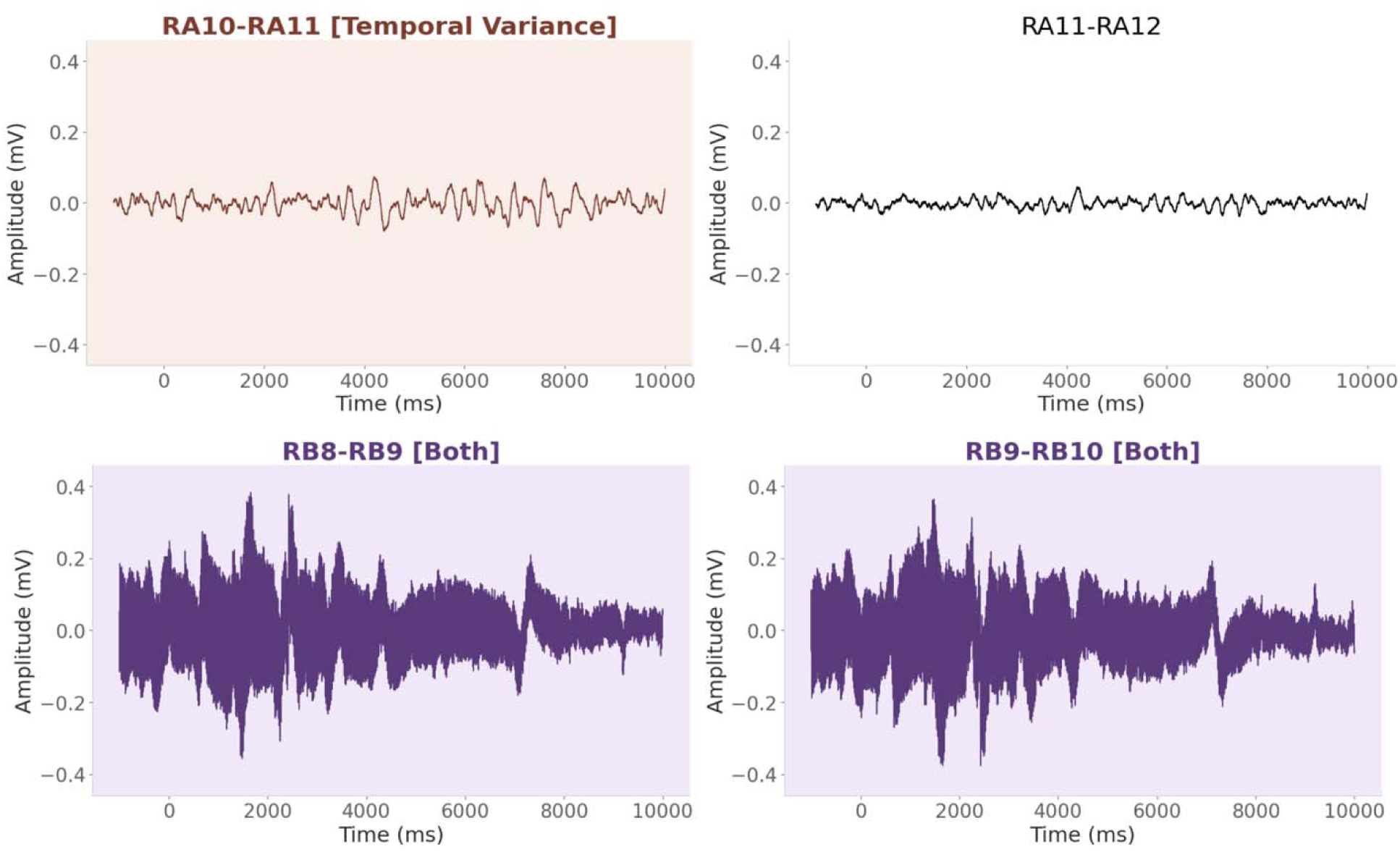
iEEG waveforms for four example bipolar contacts with artifact flagging. Mean amplitude (mV) across trials is plotted as a function of time (ms) for each bipolar channel pair (only four representative channels are shown). To identify potentially artifactual channels, two complementary variability-based metrics were computed and z-scored across channels: (1) trial-to-trial variability and (2) temporal variability. Channels exceeding |z| > 2 on both metrics are highlighted in purple (RB8-RB9 and RB9-RB10 in this example); channels flagged by temporal variance alone are shown in red (RA10-RA11); channels flagged by trial-to-trial variability alone are shown in blue (not shown in this example); and unaffected channels are shown in black. channels flagged by the automated quality-assurance metrics and the underlying electrophysiological signals during final expert review.

#### Agreement with Routine Expert Channel Annotations

Although the proposed statistical metrics are intended as quality-assurance tools rather than automated channel-rejection algorithms, it is nevertheless useful to determine whether they identify channels that experienced reviewers have independently judged to be problematic. We therefore evaluated the agreement between the automated variability metrics and routine manual channel annotations.

The comparison was performed using a retrospective multicenter iEEG dataset collected across Columbia University, Dartmouth College, Emory University, Thomas Jefferson University, Mayo Clinic, the National Institutes of Health, and the University of Texas Southwestern. The dataset comprised recordings from 65 participants (13,017 bipolar channels), including 1,185 channels that had been manually annotated as bad during routine preprocessing. These annotations were generated independently by EEG technologists and experimenters during routine data inspection and therefore do not constitute objective ground truth. Rather, they provide a pragmatic reference reflecting real-world quality-assurance decisions across laboratories and clinical sites.

To evaluate agreement between the automated statistical metrics and routine expert annotations, we treated the experts’ manual labels as the reference classification. A bipolar channel was classified as bad if either constituent contact had been labeled as artifactual during routine preprocessing. A bipolar channel was classified as good if neither constituent contact had been manually labeled as bad and the channel was not identified as belonging to the seizure-onset zone (SOZ) or containing interictal epileptiform discharges (IEDs).

Agreement between the automated and experts’ manual classifications was quantified using signal detection theory (SDT). The analysis was performed separately for trial-to-trial variability, temporal variability, their union (OR), and their intersection (AND). For each variability metric, channel z-scores were thresholded using absolute cutoffs ranging from 0 to 5 (0.5-step increments). Channels exceeding the selected threshold were automatically flagged.

At each threshold, we calculated the hit rate (HR; proportion of manually annotated bad channels correctly flagged by the automated metric), the false alarm rate (FAR; proportion of manually annotated good channels incorrectly flagged), and the sensitivity index (d′ = Z(HR) − Z(FAR)). HR, FAR, and d′ were first computed separately for each participant and then averaged across participants. One-sample *t*-tests were performed to determine whether d′ exceeded zero, with false discovery rate (FDR) correction applied across thresholds.

Both variability metrics showed significant agreement with routine expert annotations across a broad range of thresholds (Figure 10). Sensitivity (d′) remained significantly above zero for all four classification rules, indicating that channels prioritized by the automated metrics were substantially more likely to have been independently identified as problematic during manual review. Combining the two metrics using the OR criterion produced the greatest overall agreement with manual annotations, whereas the AND criterion provided a more conservative channel-selection strategy.

**Figure 10.**
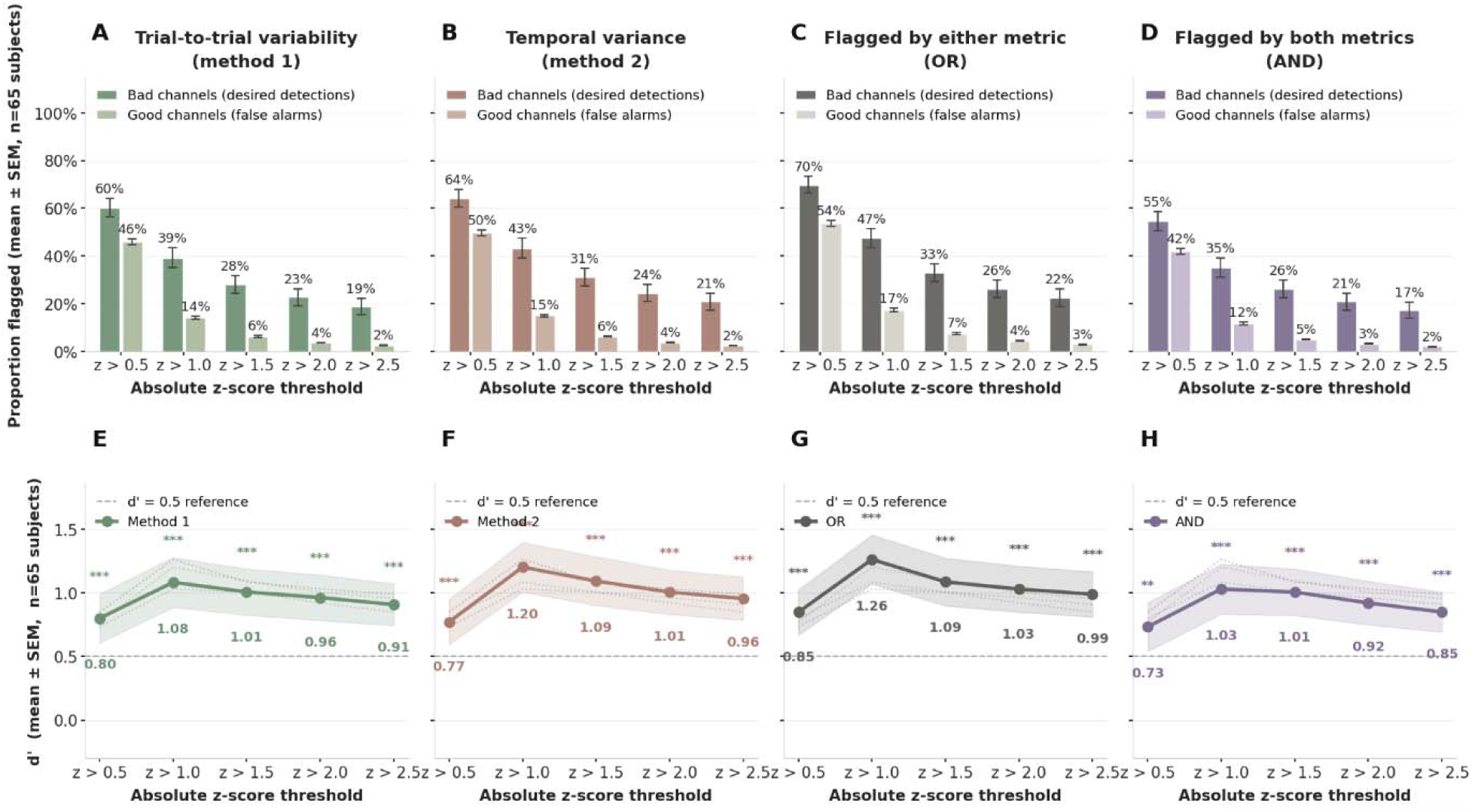
Agreement between statistical variability metrics and routine expert channel annotations. **A–D:** Mean proportion of channels flagged at each absolute z-score threshold, shown separately for manually annotated bad channels (darker color) and good channels (lighter color) across four flagging rules: trial-to-trial variability (A), temporal variability (B), either metric (OR; C), and both metrics (AND; D). Both metrics were z-scored within session prior to thresholding. Bars show mean ± SEM across subjects (n = 65). **E–H:** Sensitivity index (d′) quantifying agreement between automated channel flags and manual annotations, computed as d′ = Z(hit rate) − Z(false alarm rate). Shaded bands indicate ±1 SEM; stars denote one-sample t-tests (***p < 0.001); dashed line marks d′ = 0.5. Across thresholds, expert-annotated bad channels were flagged more frequently than good channels, resulting in positive d′ values for all methods. The OR condition showed the greatest overall agreement with manual annotations, whereas the AND condition provided the most conservative channel selection.

Because epileptiform activity can itself produce elevated signal variability independent of recording quality, SOZ and IED channels were excluded from the primary agreement analysis. Nevertheless, we quantified how frequently these channels were flagged by the proposed metrics. Both methods identified SOZ and IED channels more frequently than good channels (Figure S3), consistent with the greater signal variability associated with epileptiform activity. The proportion of flagged SOZ and IED channels remained substantially lower than that observed for manually annotated bad channels. However, when the objective is to characterize pathological neural activity, elevated statistical variability should be interpreted cautiously because the proposed metrics are designed to identify atypical signal statistics rather than distinguish recording artifacts from physiological epileptiform activity.

Importantly, these analyses are not intended to validate an automated bad-channel detector or establish channel-exclusion criteria. Instead, they demonstrate that simple statistical measures of signal variability frequently identify channels that independent reviewers had already considered problematic. Within the proposed quality-assurance framework, these metrics function as computationally efficient screening tools that prioritize channels for expert review rather than replace human judgment.

Collectively, the spectral and statistical quality-assurance measures presented in this domain provide complementary perspectives on signal integrity. Their interpretation is strengthened by integrating multiple sources of evidence - including spectral characteristics, spatial clustering of affected channels, consistency across recording sessions, and knowledge of the acquisition hardware architecture - rather than relying on any single metric in isolation.

## Discussion

Intracranial EEG (iEEG) research provides an unparalleled opportunity to study human brain function while simultaneously posing unique technical and methodological challenges. Here, we present a quality assurance framework for rapid, session-level identification of common experimental, technical, and hardware-related data-quality failures in the immediate aftermath of data acquisition. A central motivation for this framework is the recognition that iEEG research is fundamentally constrained by limited experimental opportunities. Human intracranial datasets rely on rare clinical opportunities for electrode implantation and require extensive clinical coordination, highly specialized personnel, and substantial logistical and financial investment.

Moreover, data collection often spans multiple hospital days, during which patient state, hardware configuration, recording quality, and experimental parameters may change dynamically. Under these conditions, rapid identification of data-quality issues becomes especially important. Detecting synchronization errors, stimulation-delivery failures, or malfunctioning electrodes immediately after a session may allow the research team to adjust subsequent recordings while the patient remains implanted. In contrast, delayed detection may render entire sessions unusable and result in irreversible loss of scientifically valuable data. In our experience, generating session-level reports that incorporate all domains presented herein is highly valuable for identifying subtle yet consequential sources of error, ranging from software errors in task execution to dynamic changes in signal quality caused by hardware malfunctions, incorrect referencing schemes, or synchronization failures. Many of these failure modes would not be apparent to the experimenter during data collection or through inspection of event logs alone.

The present approach focuses on four core domains, including verification of protocol fidelity, confirmation of behavioral engagement, validation of stimulation delivery, and identification of broken or contaminated channels. Each of these four domains should be adapted to the requirements of the study at hand. For example, behavioral summaries in a memory-navigation task may focus on navigation trajectories and recall accuracy, whereas language or motor paradigms may instead emphasize response timing, speech production, or movement-related measures. The modular structure of the present framework enables individual components to be tailored to the specific requirements of an experimental protocol, recording hardware configuration, and data acquisition software.

Further contributing to experimental heterogeneity in contemporary iEEG studies is the increasing complexity of direct brain stimulation protocols. Modern stimulation studies frequently manipulate frequency, amplitude, timing, anatomical target, or behavioral contingency within a single experiment. Under these conditions, accurate verification of stimulation delivery becomes essential for both scientific validity and patient safety. The present framework validates stimulation timing, frequency, and spatial specificity directly from the electrophysiological signal rather than relying exclusively on stimulation metadata logs. This distinction is important, as stimulation-command logs alone cannot guarantee that stimulation was delivered correctly at the neural recording interface. Signal-level verification provides an independent confirmation of actual stimulation delivery and enables rapid identification of timing offsets, incomplete stimulation trains, or unexpected spatial spread of stimulation artifacts.

Several limitations of the present framework should be acknowledged. First, the framework was developed and demonstrated primarily using stimulation-based memory experiments involving depth electrodes, and some visualizations or metrics may require adaptation for other paradigms or implantation strategies. Second, although the framework emphasizes rapid post-session review, interpretation of outputs still depends on domain expertise and familiarity with the experimental protocol. Automated reports can facilitate standardization and reduce oversight, but they cannot fully replace expert inspection of raw data. Third, while the pipeline flags channels with abnormal variability metrics, it does not automatically determine whether flagged channels should be excluded from downstream analyses. Such decisions depend on the specific research question, analysis strategy, and verification through visual inspection of raw signals. We intentionally designed the pipeline to provide quantitative metrics and visualizations without making automated exclusion decisions, allowing researchers to apply domain expertise and study-specific criteria.

Despite these limitations, the framework offers several practical advantages for intracranial research workflows. Automated session-level reporting can reduce preprocessing burden, facilitate onboarding and training of new personnel, improve consistency across analysts, and support reproducible documentation of preprocessing decisions. These benefits may be especially valuable for multi-site collaborations and large-scale iEEG consortia, where heterogeneity in acquisition practices and preprocessing standards can complicate data harmonization. More broadly, the increasing scale and sophistication of human intracranial EEG may require a shift toward standardized QA practices analogous to those adopted in neuroimaging and large-scale electrophysiology initiatives^8,9^. As iEEG datasets continue to expand in size, complexity, and collaborative scope, reproducible and transparent QA procedures will become increasingly important for ensuring scientific reliability and maximizing the value of these uniquely difficult-to-obtain datasets. Effective QA depends not only on electrophysiological recordings, but also on the availability of structured metadata describing behavioral events, stimulation parameters, recording configurations, and protocol expectations.

Consequently, efforts to establish common data elements and BIDS-compatible metadata standards may provide an important foundation for scalable and reproducible quality-control workflows. By leveraging standardized metadata structures, future frameworks may enable automated validation across studies, institutions, and data archives, thereby facilitating large-scale harmonization of human intracranial electrophysiology datasets. We view the present framework as an initial step toward community-wide standards for session-level quality control in human intracranial electrophysiology, which will support more reliable experimental execution, preprocessing transparency, and minimize the likelihood of irreversible data loss in human intracranial neuroscience research.

## Data and code availability statement

We illustrate the framework using de-identified intracranial EEG data collected under an experimental protocol approved by the Thomas Jefferson University Institutional Review Board (IRB #2023-2484). Example data are provided in Brain Imaging Data Structure (BIDS) format and can be accessed at https://osf.io/8yxde. The code used to generate the QA report and an example report are freely available at https://github.com/HerzLab/QC-control-paper.

## Acknowledgements

We thank Anu Chidambaram for assistance with figure preparation and graphical design.

## Notes

https://github.com/HerzLab/QC-control-paper

### Competing Interest Statement

The authors have declared no competing interest.

