## Supplemental Information for "A quality assurance framework in human intracranial electrophysiology"

### Supplementary Information

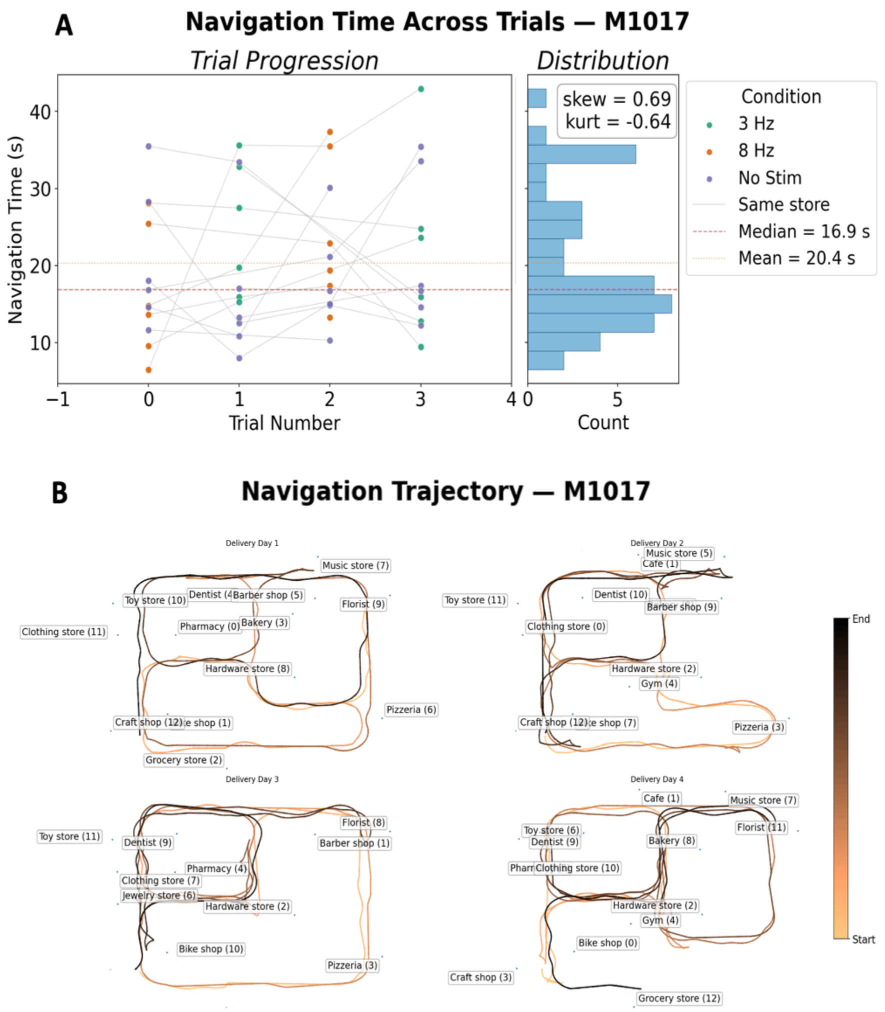


***Figure S1. Navigation time and trajectories summaries for session-level behavioral assessment. A.*** *Navigation time as a function of trial number. The right panel displays the distribution of navigation times across trials. Markedly skewed or bimodal distributions may indicate task interruption, disengagement, or technical difficulties affecting navigation.* ***B.*** *Representative spatial navigation trajectories from four trials within a session. Incomplete spatial coverage, restricted movement patterns, repetitive trajectories, or prolonged stationary periods may indicate non-random target selection, joystick malfunction, impaired navigation, or difficulty interacting with the virtual environment.*

```
LPR3-LPR4       | Power: -63.8355 dB | Z-score:  +4.06 ← Z > 2 (HIGH)
LPR4-LPR5       | Power: -63.9260 dB | Z-score:  +4.03 ← Z > 2 (HIGH)
RA1-RA2         | Power: -68.2234 dB | Z-score:  +2.85 ← Z > 2 (HIGH)
RML1-RML2       | Power: -70.4960 dB | Z-score:  +2.22 ← Z > 2 (HIGH)
LPA1-LPA2       | Power: -70.5789 dB | Z-score:  +2.20 ← Z > 2 (HIGH)
LV14-LV15       | Power: -70.7246 dB | Z-score:  +2.16 ← Z > 2 (HIGH)
RML3-RML4       | Power: -71.1527 dB | Z-score:  +2.04 ← Z > 2 (HIGH)
LPA3-LPA4       | Power: -71.7142 dB | Z-score:  +1.88
LMA3-LMA4       | Power: -71.7940 dB | Z-score:  +1.86
LPA2-LPA3       | Power: -72.1588 dB | Z-score:  +1.76
RML2-RML3       | Power: -72.2051 dB | Z-score:  +1.75
LSA2-LSA3       | Power: -72.3380 dB | Z-score:  +1.71
RML4-RML5       | Power: -72.4430 dB | Z-score:  +1.68
LBA2-LBA3       | Power: -72.6680 dB | Z-score:  +1.62
```

***Figure S2. Example ranking of channels according to the degree of line noise contamination.*** *Channels are ordered according to their mean log-transformed power within the 55–65 Hz frequency band, from highest to lowest. For readability, only the highest-ranking channels are displayed. Channels exceeding the default threshold (z > 2) are automatically flagged for review.*

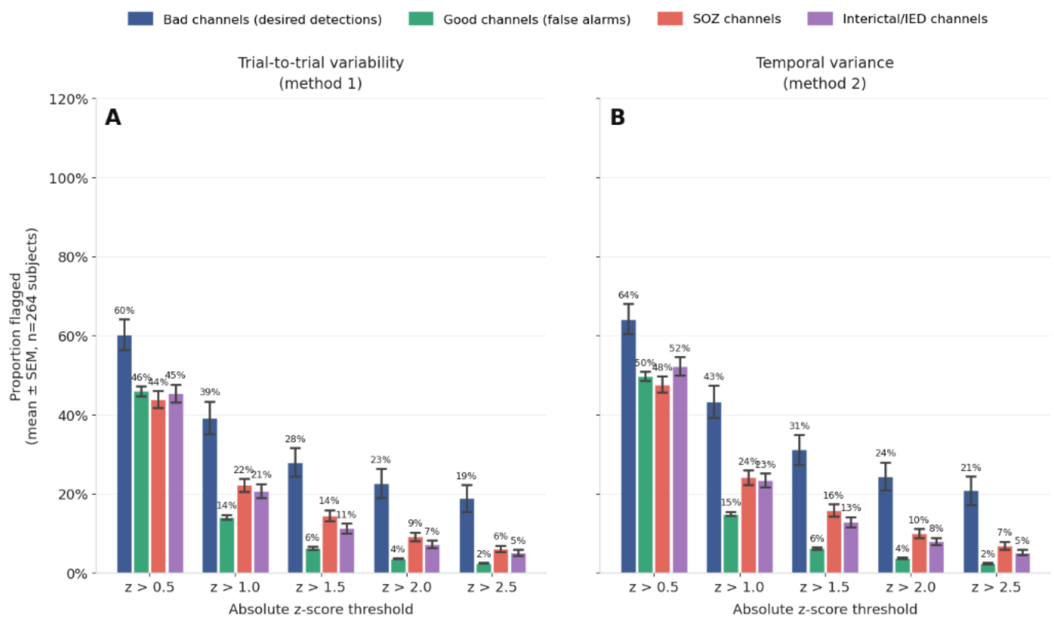


***Figure S3. Proportion of channels flagged at each z-score threshold, stratified by manual annotation category. A.*** *Trial-to-trial variability metric (Method 1).* ***B:*** *Temporal variability metric (Method 2). Channel categories reflect experts' manual annotations: bad channels (navy), good channels (green), seizure-onset zone (SOZ) channels (red), and channels containing interictal epileptiform discharges (IEDs; purple). A channel was classified as good only if it was absent from all manually annotated categories. Bars show mean ± SEM across subjects (n = 65). For both metrics, manually annotated bad channels were flagged at the highest rates across thresholds. SOZ and IED channels were flagged more frequently than good channels but less frequently than manually annotated bad channels, consistent with the elevated variance and nonstationary activity often observed in pathological neural recordings.*
